# Interpreting the CTCF-mediated sequence grammar of genome folding with AkitaV2

**DOI:** 10.1101/2024.08.01.606065

**Authors:** Paulina N. Smaruj, Fahad Kamulegeya, David R. Kelley, Geoffrey Fudenberg

## Abstract

Interphase mammalian genomes are folded in 3D with complex locus-specific patterns that impact gene regulation. CTCF (CCCTC-binding factor) is a key architectural protein that binds specific DNA sites, halts cohesin-mediated loop extrusion, and enables long-range chromatin interactions. There are hundreds of thousands of annotated CTCF-binding sites in mammalian genomes; disruptions of some result in distinct phenotypes, while others have no visible effect. Despite their importance, the determinants of which CTCF sites are necessary for genome folding and gene regulation remain unclear. Here, we update and utilize Akita, a convolutional neural network model, to extract the sequence preferences and grammar of CTCF contributing to genome folding. Our analyses of individual CTCF sites reveal four predictions: (i) only a small fraction of genomic sites are impactful, (ii) insulation strength is highly dependent on sequences flanking the core CTCF binding motif, (iii) core and flanking sequences are broadly compatible, and (iv) core and flanking nucleotides contribute largely additively to overall strength. Our analysis of collections of CTCF sites make two predictions for multi-motif grammar: (i) insulation strength depends on the number of CTCF sites within a cluster, and (ii) pattern formation is governed by the orientation and spacing of these sites, rather than any inherent specialization of the CTCF motifs themselves. In sum, we present a framework for using neural network models to probe the sequences instructing genome folding and provide a number of predictions to guide future experimental inquiries.

**Author Summary:** Mammalian genomes are spatially organized in 3D with profound consequences for all processes involving DNA. CTCF is a key genome organizer, recognizing numerous sites and creating a variety of contact patterns across the genome. Despite the importance of CTCF, the sequence determinants and grammar of how individual sites collectively instruct genome folding remain unclear. This work leverages the ability of Akita, a deep neural network, to make high-throughput predictions for genome folding after DNA sequence perturbations. Using Akita, we make several experimentally testable predictions. First, only a minority of annotated sites individually impact folding, and flanking DNA sequences greatly modulate their impact. Second, multiple sites together influence folding based on their number, orientation, and spacing. In sum, we provide a roadmap for interpreting neural networks to better understand genome folding and important considerations for the design of experiments.

## Introduction

Mammalian genomes contain thousands of CTCF sites, which collectively instruct locus-specific 3D genome folding [1,2]. These sites influence genome architecture through CTCF’s ability to halt loop extrusion by the cohesin complex. The global importance of CTCF and cohesin are evident by the loss of topologically associated domains (TADs) and dots (also termed loops) upon the depletion of either factor [3,4]. Despite this, the impact and function of individual CTCF sites remain less understood.

Genome folding is thought to either insulate or facilitate physical interactions between regulatory sequences, thereby modulating enhancer-promoter communication [5]. Localized disruptions at specific CTCF sites have displayed clear changes to genome folding, gene expression, and development. Disrupted folding at the EPHA4 locus is linked to limb phenotypes [6], and changes to individual CTCF sites at the same locus are associated with index finger phenotypes [7]. Targeted deletion of CTCF sites at the Kallikrein locus led to coordinated gene activity associated with prostate cancer [8]. Nevertheless, disruptions to many CTCF sites have little or no impact; for example, [9] observed that the fusion of two adjacent TADs after deletion of multiple CTCF sites at the Kcnj2/Sox9 locus resulted in no detectable phenotype. An understanding of when sequence perturbations at TAD boundaries or CTCF sites would actually disrupt genome folding is a prerequisite for understanding downstream impacts of genome folding on gene regulation.

Which CTCF-binding sites are essential for genome folding, and why are they particularly crucial? Even the highest-resolution mammalian Hi-C and Micro-C assays are challenging to analyze below a resolution of 1kb, much larger than the approximately 20bp-long motifs recognized by CTCF. Additionally, individual pixels at dots or boundaries often contain more than one CTCF motif. How do collections of CTCF sites instruct the formation of features on Hi-C maps? It has been shown that TAD boundaries are enriched with divergent CTCF sites [10], while dot anchors are enriched with convergent CTCF sites [11]. Altering the orientation of CTCF sites alters local patterns of genome folding [6,12–14]. While experimental techniques can now quantify the impacts of specific DNA sequence perturbations on genome folding and gene expression [15–18], testing throughput is currently limited to dozens of sites in a given genome context.

Computational approaches enable high-throughput screening of hundreds of thousands of flexibly-designed sequences *in silico*. Deep neural network models can now rapidly generate state-of-the-art predictions of genome folding from DNA sequences [19–22]. Using Akita, one of these models, we found that roughly 40% of nucleotides predicted to strongly impact local genome folding are located within the 100 nucleotides flanking CTCF motifs [19]. However, these impacts remained poorly characterized. Methods to interpret neural networks trained on genomic data offer promising insight into sequence-based mechanisms [23]. Recent applications included learning the motif syntax and grammar of transcription factors, such as Nanog [24], OSK, and AP-1 [25]. Together, this argues that further investigation of CTCF flanking sequences by interpreting trained neural networks can provide a deeper understanding of how DNA sequence determines 3D genome folding.

Here we update the Akita framework to jointly leverage mouse and human genome folding data and use this model to quantify the sequence contributions to locus-specific 3D folding (**Fig 1**). We screened millions of sequences *in silico* and quantified their predicted impact on genome folding. We observed a surprisingly low correlation between predicted disruption to chromosome structure upon mutating a CTCF site and CTCF ChIP-seq data, yet a high correlation with the frequency of binding measured by single-molecule footprinting experiments. By inserting thousands of strong CTCF motifs into background sequences and assessing their impact, we identified a critical role of flanking sequences for determining the most significant CTCF sites for genome folding. We find that pairs of mutations within CTCF motifs are largely additive and that the strength of CTCF site clusters depends on the number of sites, their orientation, and spatial arrangement. Finally, we find that CTCF motifs instruct genome folding in a feature-agnostic manner rather than preferentially forming either dots or TAD boundaries. Collectively these results deepen our understanding of the sequence preferences and grammar through which CTCF contributes to genome folding.

**Figure 1.**
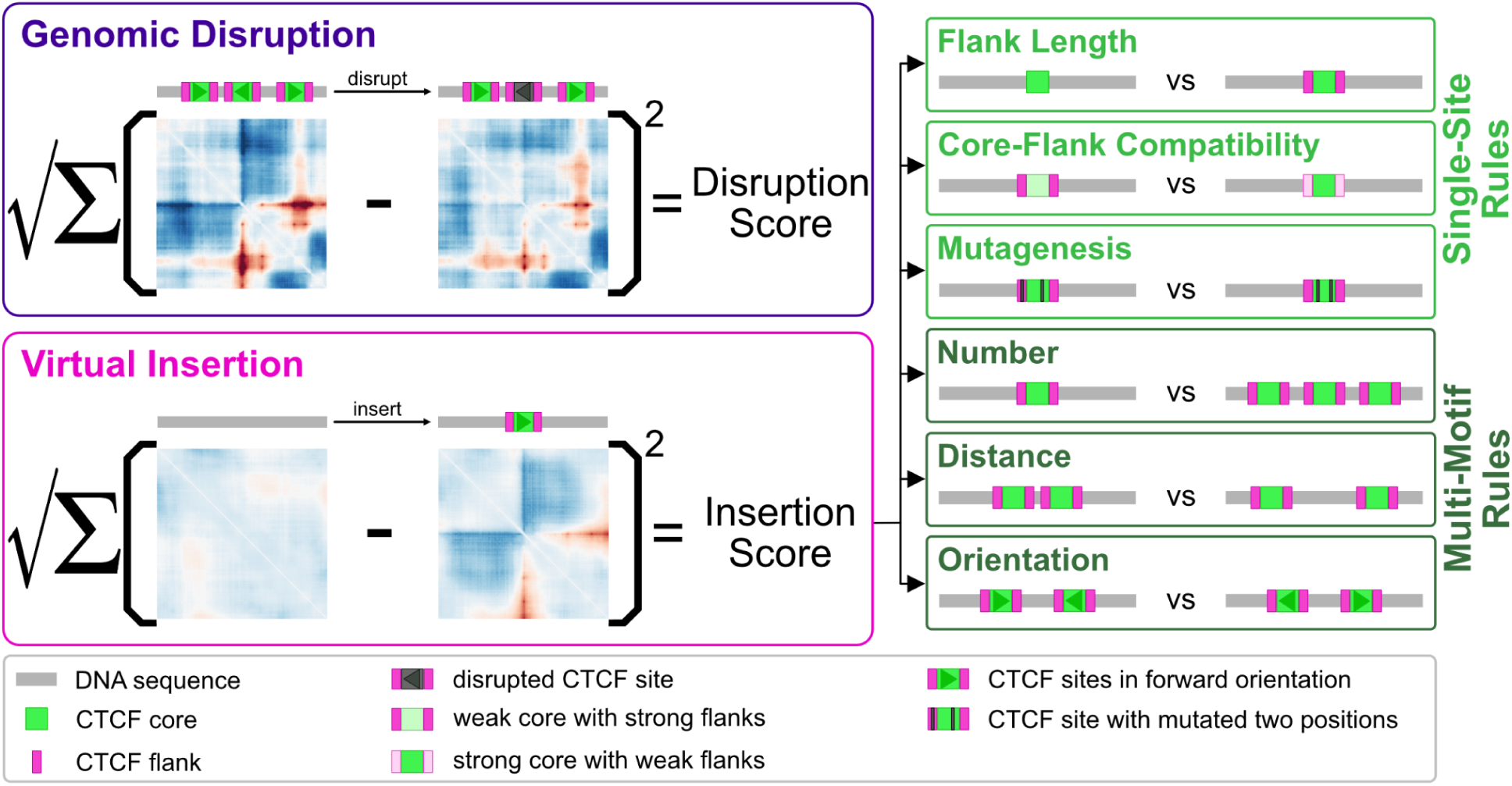
Utilizing AkitaV2 to extract CTCF-directed sequence preferences and grammar. Conceptual summary of the analyses performed in this study. *Left:* two main approaches: genomic disruption and virtual insertion. Genomic disruption involves permuting the nucleotides of a CTCF site within its genomic context, while virtual insertion entails inserting a CTCF site into a feature-less background sequence. Sequences and motifs are shown as cartoon sequences with illustrative predicted maps below. *Right:* six types of virtual insertion experiments reveal distinct aspects of CTCF-site grammar. Three experiments tested single-site rules: (1) the impact of flanking sequences, (2) the compatibility between core motifs and their flanking sequences, and (3) nucleotide level mutagenesis. Three experiments tested multi-motif grammar: (4) varying numbers of CTCF sites within a cluster, (5) varying spacing between sites, and (6) orientation. Cartoon sequences represent the parameters tested in these experiments.

## Results

### Cross species model

Based on the observation that cross-species training can improve neural network predictions for sequence-to-profile models [26], we employed a similar approach to update Akita. Following the approach of the cross-species Basenji model [26], we trained an ensemble of eight models, each with a distinct held-out subset of the two genomes. Each joint model was trained on 6 mouse and 5 human high-quality Hi-C and Micro-C datasets as targets (**Fig S1A**) with a slightly increased input sequence length (now 1.3Mb for AkitaV2, up from 1Mb). Trained models predict log2 observed/expected contact frequency at 2048bp resolution for any given input sequence. We observed a modest performance increase for AkitaV2 measured as the correlation of predictions with held-out test data (Pearson R=0.66 vs. 0.61 previously, **Fig S1B-E**).

To quantify the influence of short DNA sequences on genome folding, we defined a disruption score as the square root of the sum of squared differences between predicted maps before and after local sequence perturbations (**Fig 1**), as previously used to interpret Akita’s predictions [19,27]. This disruption score is sensitive to gain or loss of boundaries, as well as changes in TAD substructures [28,29]. We leveraged the ensemble of models to validate sequence perturbation approaches at CTCF sites by their cross-model stability. For AkitaV2, we found that masking (i.e. replacing nucleotides with zeros) was not consistent across models, while disruption by random permutation displayed several favorable properties. Predicted disruption scores by permutation are: (i) highly correlated across random permutations for any given CTCF site; (ii) consistent across models; (iii) robust regardless of whether the perturbed site is in the center or shifted by up to 10kb in the input DNA; (iv) preserved for the reverse-complement (**Fig S2A-D**). These observations argue that disruption by permutation is a robust strategy for extracting predicted impacts of various CTCF sites with Akita. Since disruption scores were highly correlated between human and mouse model outputs (Pearson R = 0.96, **Fig S2E**), we focused on predictions from the mouse model for subsequent analyses.

### CTCF ChIP-seq provides poor prioritization for impactful sites as assessed by genomic disruptions

Using our updated model and validated perturbation approach, we analyzed the relationship between disruption scores and mouse epigenomic features thought to relate to genome folding (**Fig 2A**). Despite targeting sequences directly underlying CTCF sites, we found only a moderate correlation with CTCF ChIP-seq (**Fig 2B**). This aligns with recent reports that insulator activity displays little or no relationship with CTCF ChIP-seq signal [15,17]. In contrast, we observe a high correlation with cohesin (RAD21) ChIP-seq and with CTCF site occupancy as measured by single-molecule footprinting (SMF) [30].

**Figure 2.**
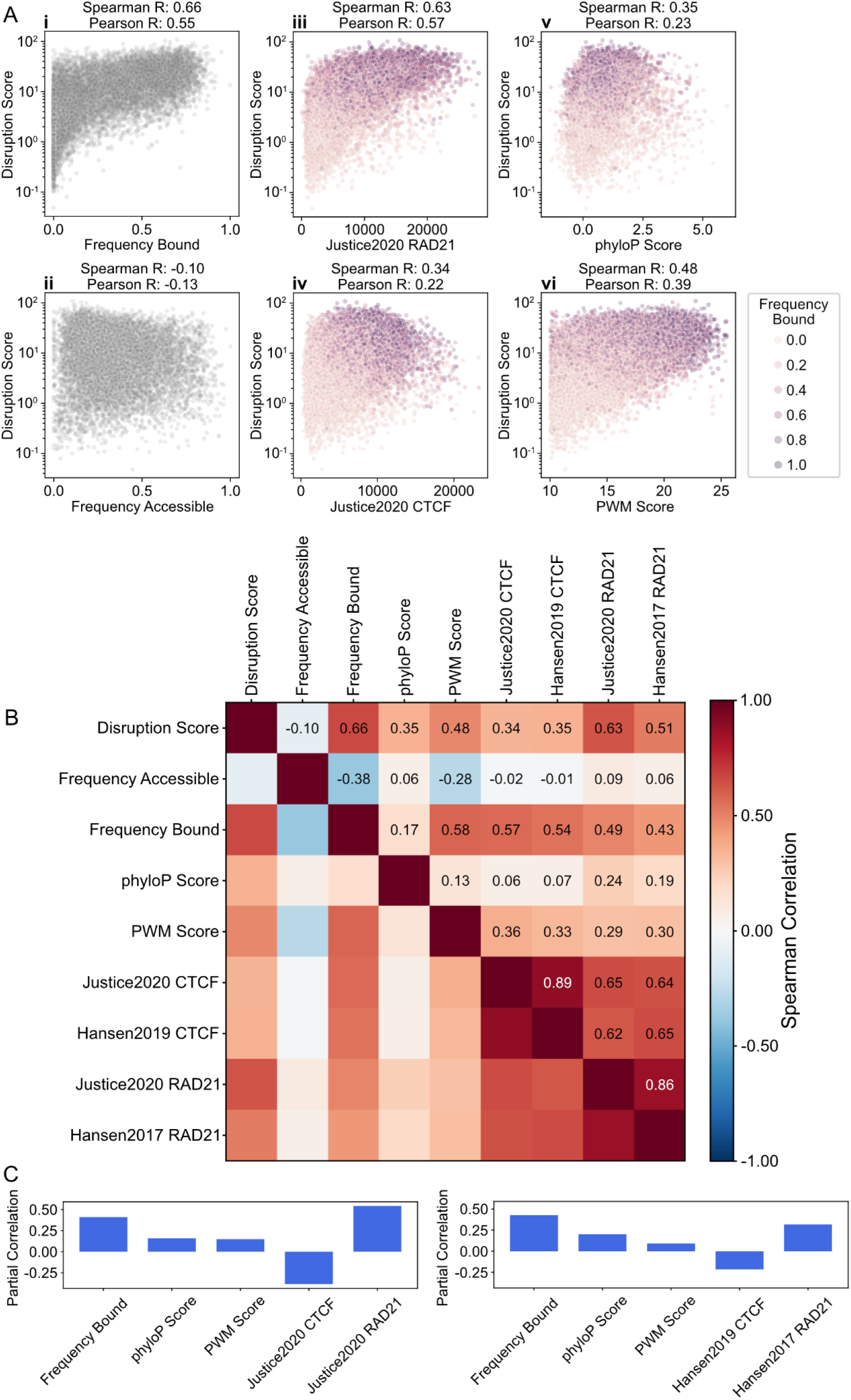
Disruption scores highlight impactful epigenomic features. **A)** Scatterplots showing disruption scores vs. genomic features at n=9,991 autosomal CTCF sites, profiled with single-molecule footprinting (SMF) [30] which categorizes sites as: bound, nucleosome occupied, or accessible. The first column displays disruption score vs. (i) frequency of being bound or (ii) accessible. The other subplots show disruption scores vs. following genomic features: (iii) Rad21 ChIP-seq signal [53], (iv) CTCF ChIP-seq signal, (v) conservation score (phyloP), and (vi) PWM score, with dots colored by their SMF bound frequency. ChIP-seq signal is quantified as the sum in a ±100bp window around the motif position. **B)** Matrix of pairwise correlations between disruption scores and genomic features of n=9,991 autosomal CTCF sites. **C)** Partial correlation coefficients between disruption scores and subsets of genomic features from panel B, adjusting for mutual influences among these features. Partial correlations computed controlling for CTCF and cohesin ChIP-seq either from [53] (left) or [54,55] (right) are similar qualitatively and quantitatively.

To quantify the relative importance of each epigenomic feature, we computed their partial correlations with our disruption scores. We observed CTCF SMF occupancy and cohesin ChIP-seq exhibited large positive partial correlations (**Fig 2C**), indicating they provide orthogonal sources of information for predicting disruptions. CTCF ChIP-seq, in contrast, displayed a negative partial correlation when accounting for the information from RAD21 ChIP-seq and SMF CTCF occupancy. This argues that CTCF ChIP-seq provides redundant information once SMF occupancy is accounted for, and highlights the benefits of new technologies for better understanding 3D genome organization.

### Virtual insertions probe CTCF influences independently of genomic context

One possibility for the low observed correlations between epigenomic signals and our CTCF disruption scores were the distinct genomic contexts of each site. For example, redundant TAD boundaries could mask the effects of CTCF perturbations in their native genomic context [6,7]. To quantify CTCF site impacts independent of their genomic context (**Fig 3A**), we developed a virtual insertion screening approach where a large set of genomic CTCF sites are inserted into neutral, largely featureless, background sequences, similar to previous approaches [24,27]. We then computed an “insertion score” for each sequence as the sum of squared differences for predicted maps before versus after the insertion (**Fig 1**).

**Figure 3.**
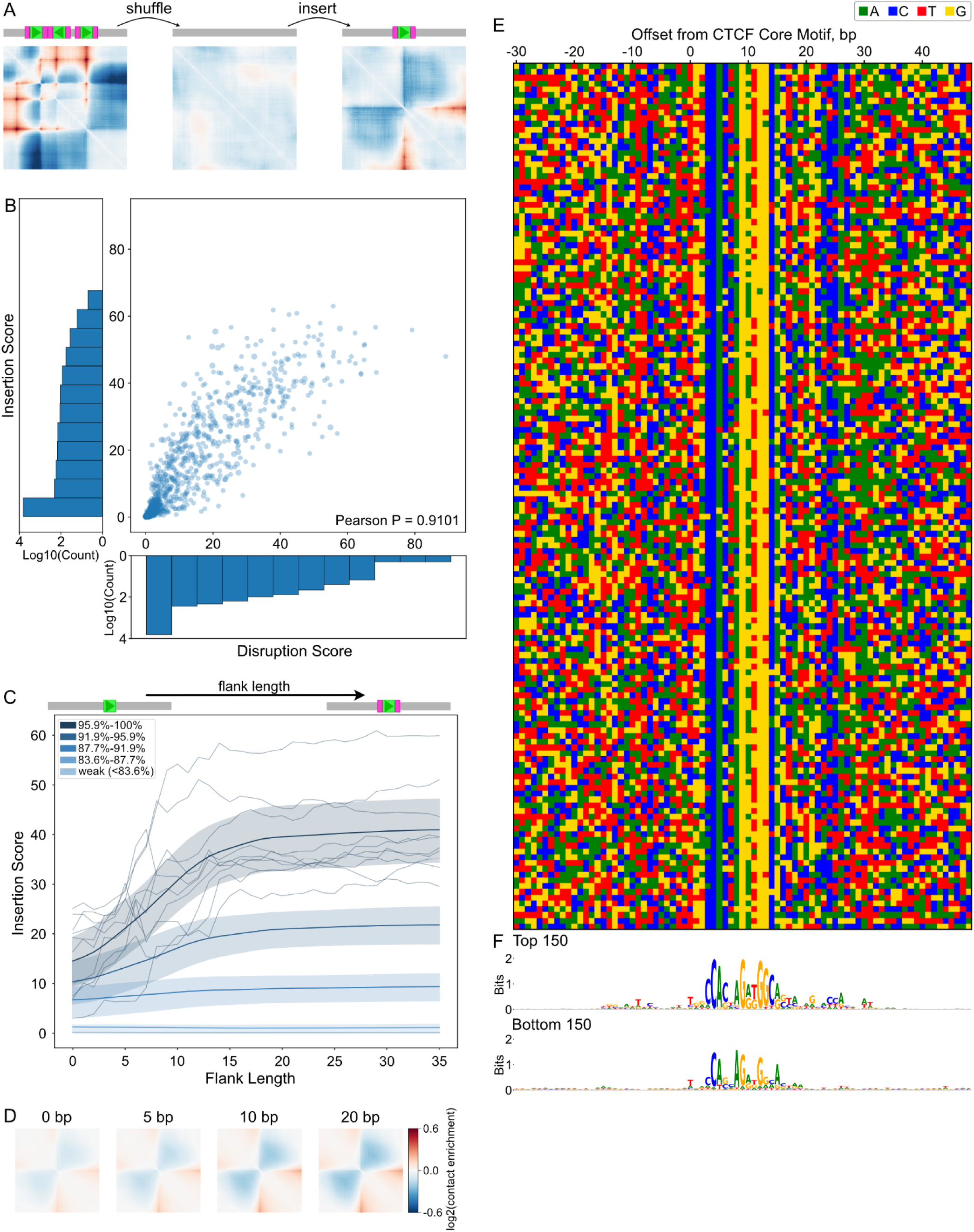
A virtual insertion strategy reveals the impact of flanking sequences. **A)** Virtual insertion strategy assesses individual CTCF site impacts. We generated background sequences by shuffling genomic sequences such that they produce mostly-featureless predicted maps. A CTCF site (green box) along with its flanking sequences (pink box) is then inserted into these background sequences (in gray). Using the sequence with an insertion as input, we generated predicted maps and quantified the impact as an insertion score. **B)** Scatterplot of insertion versus disruption scores, for n=7,560 CTCF sites (PearsonR > 0.91). Sites were obtained by intersecting JASPAR motifs with mESC boundaries and filtering for lack of overlap with repetitive elements within +/- 20bp or other CTCF sites within +/- 60bp. Scores were averaged across all six mouse outputs (i.e. cell types) and all eight models. Insertion scores were additionally averaged over ten background sequences. Histograms show log density along each scatterplot axis, as the majority of sites exhibit both low insertion and disruption scores. Given this, for further analysis we selected the 1250 sites with the highest disruption scores and chose an additional 250 sites randomly from the remaining pool. **C)** Flanking sequence length versus insertion score for the analysis set of n=1,500 CTCF sites. Flanking sequence was varied from 0bp (19bp core CTCF motif only) up to 35bp, depicted as cartoons above the plot. Genomic flanking sequences were symmetrically extracted around each CTCF site. For visualization, sites were divided into five groups based on their insertion score with 30bp flanks. Smoothed lines show the mean for each group, and shaded bands show the 25th to 75th percentiles of motifs within the group. To illustrate the variability among sites, we show 10 sites chosen randomly from the strongest group as navy lines. **D)** Predicted contact maps illustrate the impact of increasing flanking sequence lengths for a strong CTCF site. Sequence of inserted CTCF site and flanks obtained from chr15:101,984,508-101,984,527 in the mouse genome. **E)** Heatmap of nucleotide composition around 150 strong CTCF sites (±30bp). Rows ordered by insertion score. **F)** Sequence logos for the sequences with top 150 and bottom 150 insertion scores highlight core motifs and flanking preferences.

To generate neutral background sequences, we found that shuffling by 8mers led to relatively featureless maps (**Methods, Fig S3A,B**). Shuffling greatly reduced the variance of insertion scores while leaving the mean unchanged (**Fig S3C**), indicating that computationally inexpensive shuffling can reduce the number of neural network predictions needed to make a reliable estimate of a CTCF site’s impact. We confirmed the robustness of our virtual insertion strategy by using the ensemble of models provided by AkitaV2. We found that insertion scores are highly consistent across backgrounds (Pearson R > 0.99, **Fig S3D**) and across different models (Pearson R > 0.96, **Fig S3E**).

To obtain CTCF sites that include those with boundary-forming potential, we extracted positions of boundaries from mESC Hi-C data [31] and overlapped these with CTCF sites from JASPAR [32]. Recognizing that CTCF motifs can be present yet not bound within repeat elements, like B2 SINEs in the mouse genome [33], we filtered out sites that overlapped repetitive elements. To prevent the inclusion of additional CTCF sites within extended flanking sequences we filtered out CTCF sites located less than 60bp from another site. By removing these potential confounders, we sharpened our focus on individual CTCF site impacts. We found a high correlation (Pearson R > 0.99, **Fig S3F**) between human and mouse predictions of mouse CTCF virtual insertions, demonstrating that a cross-species model with high predictive power utilized very similar sequence patterns at the motif level.

We observed a strong correlation between virtual insertion scores and genomic disruption scores (Pearson R > 0.91) for the filtered CTCF sites (**Fig 3B**). Despite coming from regions specifying TAD boundaries, the vast majority of CTCF sites exhibited low disruption and insertion scores, and only a small fraction has a considerable predicted impact on genome folding. The great differences between CTCF sites that we predict helps understand experimental observations where deleting different CTCF sites, even within the same boundary, led to distinct outcomes for genome folding and gene expression [7,9,34]. Given the great number of weak sites, for deeper analysis we focused on a set of 1500 sites, including 1250 sites with the highest scores and 250 sites picked randomly from the remaining pool.

### Flanking sequences modulate the influence of CTCF sites on genome folding

We next tested how flanking sequences around CTCF sites influenced predicted insertion scores. We defined core motifs as the 19bp-long sequences from JASPAR (MA0139.1), and flanking sequences as the genomic sequences up- and downstream of the core. We inserted individual CTCF sites into neutral background sequences, incrementally extending the flanking genomic sequence around the core motif. We observed that average insertion scores rose sharply with increased flank length up to about 15bp before stabilizing (**Fig 3C,D**). Our finding is consistent with experimental observations that highlight the importance of flanking sequences for transcriptional insulation [15,17], binding [35,36], and accessibility [37]. We repeated the flanking sequence insertions with two copies of each CTCF site and observed a similar trend for all four possible orientations (**Fig S4B-D**). This argues that the impact of flanking sequence on an individual CTCF site strength is independent of other nearby sites.

Recognizing the strong contribution of flanking sequences, we searched for sequence preferences within these regions. We generated sequence logos for CTCF sites with the highest and lowest insertion scores. In the flanking regions of motifs with low insertion scores, we found little sequence preference (**Fig 3F**). In contrast, flanking regions around strong CTCF sites displayed sequence preferences 2 to 13bp downstream and −15 to −7bp upstream of the core motif, though these were more subtle than the core motif itself (**Fig 3E**). Influential flanking sequences for genome folding, as predicted by AkitaV2, aligned well with previously documented CTCF binding preferences upstream and/or downstream of the core motif derived from ChIP-seq, MNase-seq, and ChIP-exo [35–40]. The sequence preferences we observed differed from those reported by [15], possibly due to a limited number of experimentally tested sites. Our virtual insertion and genomic disruption scores produced largely congruent motif logos (**Fig S4A**). Consistent with [27], we found that thymine at the 8th and 12th positions was elevated for the strongest sites relative to the consensus motif.

We hypothesized that subtle sequence preferences might reflect an average over distinct binding modes. We explored multiple methods to order and cluster flanking sequences around strong sites, including by: overall insertion scores (**Fig 3E**), the Hamming distance between upstream, downstream or combined upstream and downstream sequences. None of these revealed clear clusters of sequence preferences (**Fig S5**). We also performed motif enrichment and *de novo* motif discovery in the flanking sequences with Homer [41]. Neither approach yielded a prevalent flanking sequence motif; the most prevalent *de novo* motif occurred in 20% and the most prevalent known motif in 10% of flanks. Together, our results point towards flanking sequences around CTCF having an important impact on genome folding, albeit one that is not readily characterized as a single position weight matrix.

### Core motifs and flanking sequences are broadly compatible

Given the absence of strong motifs in the flanking sequences themselves, we investigated whether flanking sequence impacts are contingent on their associated core motif sequences. We assessed predicted core-vs-flank compatibility using 300 CTCF sites classified as strong, medium, or weak based on their overall insertion scores. We inserted all possible core-flank combinations into background sequences and assessed the strength of the resulting maps (**Fig 4A**). If compatibility was an important factor, insertion scores for cognate core-flank pairs from the genome would be stronger than synthetic combinations. We found no evident core-flank compatibility: there was no evidence of higher scores for genomic cognate pairs (i.e. no strong diagonal in the pairwise matrix, **Fig 4D**), and no clear deviation of the distribution of scores for cognate versus synthetic pairs (**Fig 4B**). Weak cores paired with strong flanks yielded relatively weak sites, while strong cores with weak flanks performed almost on par with medium cores paired with medium flanks. These predictions agree with experiments that quantified the impact of core and flanking sequence from strong versus weak sites on transcriptional insulation [15]. We performed singular value decomposition (SVD) to factorize the matrix of pairwise core-flank combinations into a set of vectors that described the influence of each core or flanking sequence. We found that the product of the first SVD factors for cores and flanks nicely approximated the insertion score for the corresponding core-flank combination (**Fig 4C,E**). This indicates that the rules learned by AkitaV2 for combining core and flanking sequences are largely multiplicative without strict constraints on their compatibility.

**Figure 4.**
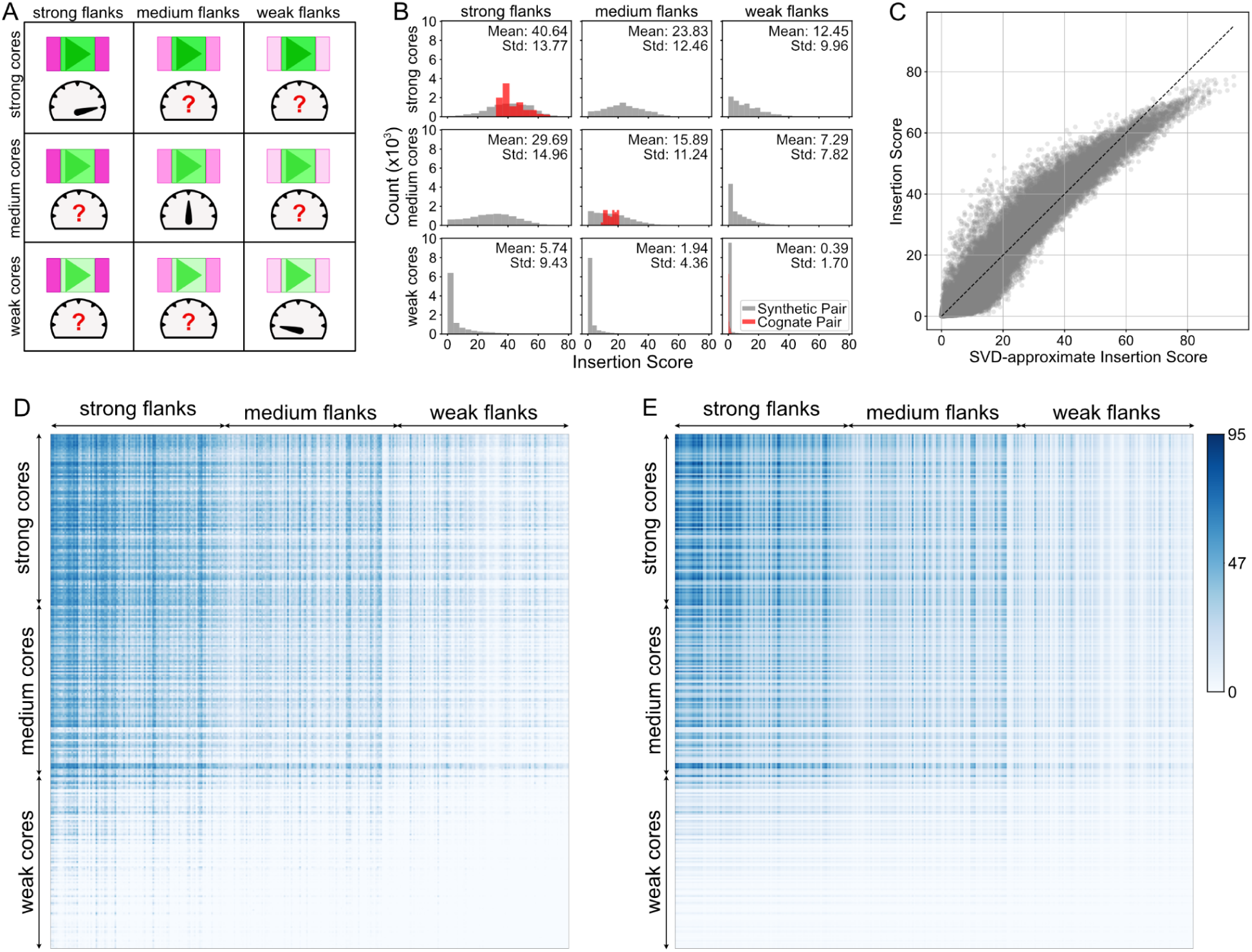
CTCF core and flanking sequences are broadly compatible. **A)** Illustration of the test for compatibility between core and flanking sequences by assessing all possible combinations of flanks and cores classified into three strength groups. Each row represents a distinct 19bp core motif sequence and each column represents a distinct pair of 30bp flanking sequences adjacent to the core motif. **B)** Distributions of insertion scores for pairs of core and flanking sequences around 100 strong, 100 medium, and 100 weak CTCF sites. Each histogram shows 10,000 (100^2^) combinations. Sites were classified as strong, medium, and weak based on their combination of core and flanking sequence seen in the mouse genome. Distributions for original genomic core-flank pairs shown in red (with count scaled by 100), synthetic combinations shown in gray. **C)** Scatterplot of insertion scores (panel D) versus approximate values obtained through SVD (panel E). Their high correspondence indicates that predicted strengths are largely multiplicative and core and flanking sequences are largely compatible. **D)** Heatmap of insertion score for 300 CTCF core and 300 flanking sequence pairs. Each row corresponds to a core CTCF motif, while each column represents a different flanking sequence. Rows and columns are ordered by the insertion score of the core-flank combination that occurs in the genome (i.e. by values along the diagonal). **E)** Heatmap of approximate insertion strength obtained via SVD for 300 CTCF core and 300 flanking sequence pairs. Rows and columns are ordered as panel D.

### Individual nucleotides within a motif contribute additively to genome folding patterns

After observing no strict compatibility requirements at the level of entire core and flanking sequences, we next explored compatibility at the nucleotide level. Given that each zinc finger domain of CTCF is thought to recognize a triplet of DNA base pairs [42], we hypothesized that pairs of mutations within the same triplet might be more detrimental than those spanning different triplets. This would create blocks of low mutation scores along the diagonal of a pairwise mutation matrix. We conducted pairwise mutagenesis on 100 strong CTCF sites that displayed the highest insertion scores. Contrary to our initial hypothesis of an epistatic effect within zinc finger triplets, we observed no clusters of heightened pairwise impacts along the diagonal (**Fig 5A**). To better understand pairwise impacts, we tested if their impacts deviated from simple additivity. To generate the additive expectation, we performed saturation mutagenesis for single mutations (**Fig 5B,C**) and summed together the scores for pairs of mutations. For weak mutations, the pairwise mutational impact was congruent with the additive expectation. After a strong mutation, however, the impact of additional mutations saturated and diverged from the additive expectation (**Fig 5D**), suggesting negative epistasis [43,44]. We hypothesize that this arises because the strongest mutations abolish CTCF binding and hence ability to impact genome folding.

**Figure 5.**
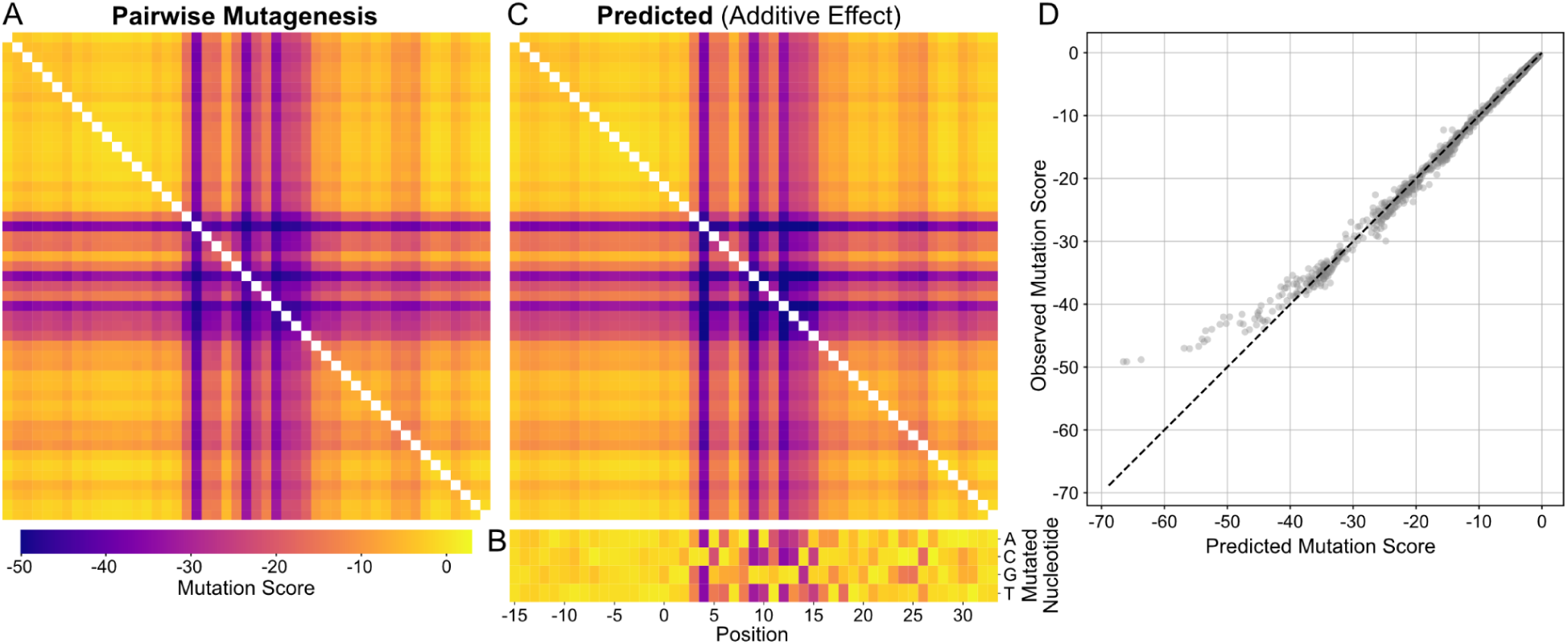
Pairwise nucleotide dependencies are largely additive in core CTCF motifs and their flanking sequences. **A)** Pairwise mutagenesis. Mutation score for pairs of mutations in the 19bp core motifs +/- 15bp flanks for 100 strong CTCF sites. Mutation score is calculated as the difference between insertion scores for the mutant versus the unperturbed sequence. The heatmap shows the average mutation score for each pair of positions. **B)** Single-nucleotide saturation mutagenesis of the 19bp core motifs +/- 15bp flanks for the same sequences in A. The heatmap presents the average over all CTCF sites for each possible substitution. **C)** Predicted additive impact of pairwise mutagenesis. The predicted additive pairwise impact is the sum of the average single-nucleotide impacts (panel B). Note the shared color scale across panels A-C. **D)** Scatterplot of predicted additive and observed pairwise mutagenesis effects from panels A,B. For pairs of weak mutations, impacts are largely additive (up to mutation scores of −40). Higher impact mutations (i.e. more negative mutation scores) appear to saturate and diverge from this linear trend.

### The positioning and orientation of multiple CTCF sites specifies a broad range of folding patterns

After quantifying the sequence preferences of individual motifs, we next turned to deciphering the multi-motif CTCF grammar by systematically varying their: (i) number, (ii) spacing, and (iii) orientation.

We found that larger numbers of inserted motifs produced correspondingly larger predicted insertion scores (**Fig 6A**). This aligns with experimental observations that insulation strength increases with more inserted CTCF sites [15] and that stronger TAD boundaries contain greater numbers of CTCF sites [45,46]. We found that dosage-dependent insulation requires strong CTCF sites, as the insertion scores for weak sites remained low. Similarly, experiments show that tandem arrays of CTCF sites from non-boundary regions do not function as insulators [15]. Our observation indicates that clusters of strong CTCF sites could be used to make stronger TAD boundaries.

**Figure 6.**
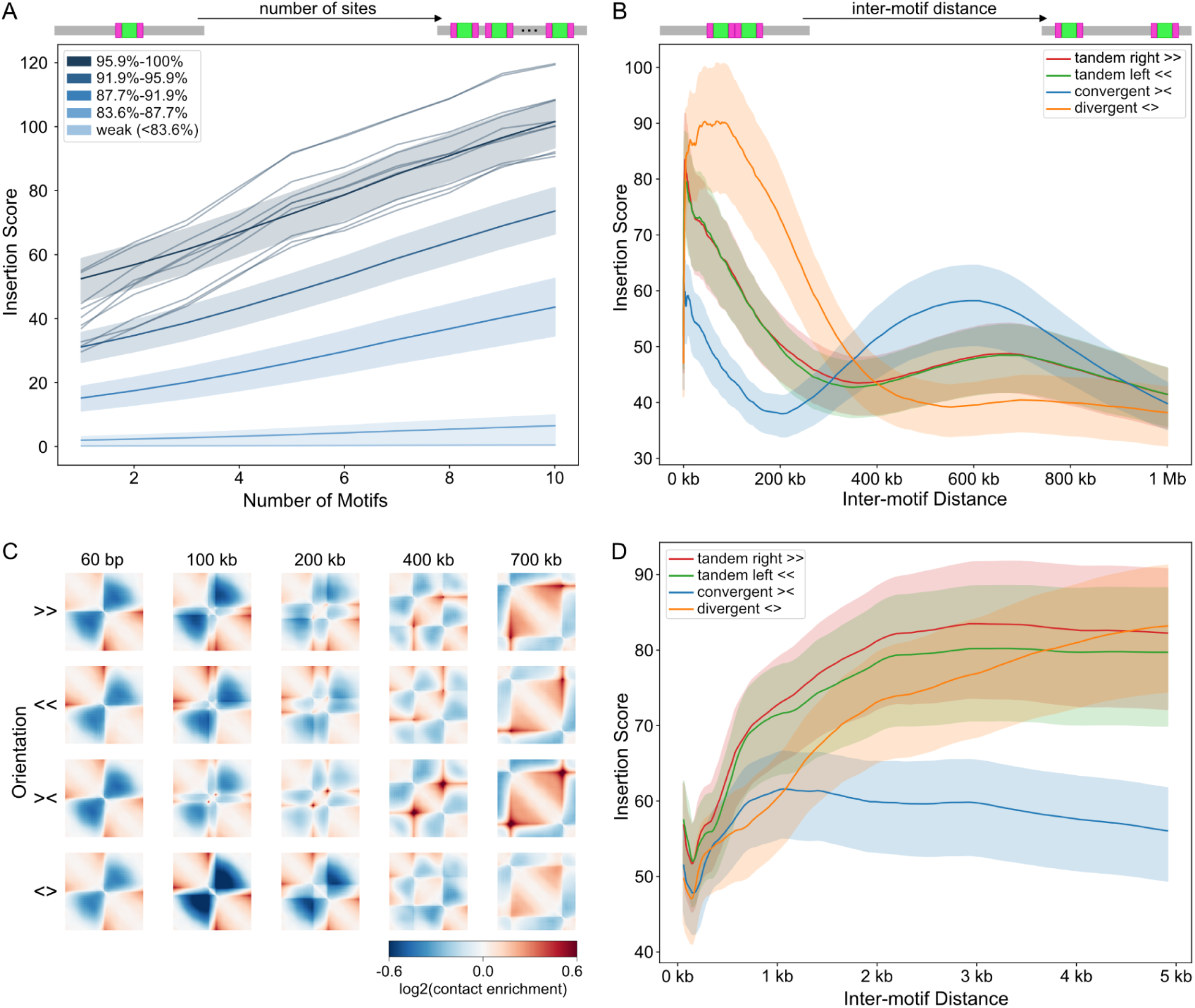
CTCF grammar depends on motif number and spacing. **A)** Insertion score versus number of inserted CTCF sites. Averages over five groups of n=1,500 CTCF sites plotted as in Fig. 3C. Shaded areas indicate 25-75th percentile for each group. Variability among sites is highlighted using 10 randomly chosen sites from the strongest group (dark navy lines). All sites inserted in a rightward orientation with 30bp flanks and 180bp spacing between cores. Note that with this spacing, 10 inserted motifs constitutes about 2kb or one bin. **B)** Insertion score as a function of motif spacing for four possible motif pair orientations for 300 CTCF sites (the strongest 20% from A), also with 30bp flanks. Average across sites shown for each orientation, with variability indicated by 25-75 percentile bands. **C)** Predicted log-transformed observed/expected contact frequency maps by CTCF pair orientation and spacing for insertion of a representative CTCF site (sequence from chr2:93,199,043-93,199,062). **D)** As in B), zooming into the 0-5kb region.

We explored how genomic context modulates CTCF grammar by inserting hundreds of strong pairs of sites at varying genomic distances and in four distinct orientations. The most notable differences in predicted maps occurred between convergent and divergent orientations. Divergent sites exhibited their highest impacts at a relatively short distance (∼70kb) before declining, whereas convergent sites displayed high impacts at two distances, including at a much greater secondary distance (∼600kb) (**Fig 6B**). Visual inspection indicated that this second maxima corresponded to dot patterns in the predicted maps (**Fig 6C**). We also observed an initial dip in insertion score at an inter-motif distance of ∼170bp, consistent across all four orientations (**Fig 6D**). Interestingly, this is close to the nucleosome repeat length estimated from MNase-seq [38], and strong CTCF sites are often flanked by arrays of phased nucleosomes [47,48]. The profiles for tandem left and tandem right sites are closely aligned, confirming the model’s strand independence. These observations demonstrate the complexity of genome folding, as even two CTCF sites can generate a diversity of features within predicted maps (**Fig 6C**).

Motivated by the distinct maps for convergent versus divergent pairs of CTCF sites, we tested whether there are categories of CTCF sites that exhibit feature specialization. To test this hypothesis, we designed an *in silico* screen using pairs of motifs positioned in two distinct scenarios: (i) convergent motifs (><) spaced 400kb apart, to test their dot-formation ability; (ii) divergent motifs (<>) with a smaller spacing of 180bp, to test their boundary-formation ability (**Fig 7A**). From predicted maps, boundary strength was estimated using the insertion score, while dot strength was assessed as the enrichment between dot anchors versus surrounding regions (**Methods**). Collectively, CTCF sites followed a consistent trend for dot versus boundary strength (**Fig 7B,C**), arguing against the feature specialization hypothesis. We also found that insertion scores from boundary scenarios were the same magnitude as those for dot scenarios (**Fig S7A**), suggesting that while the number of sites influences the overall strength of the predicted map, the spacing and orientation determine which features are visible. To rule out bias from our selection of inserted sites, we repeated the analysis with CTCF sites specifically overlapping dot anchors, as identified by *MUSTACHE* [49]. We found a largely similar trend in terms of dot versus boundary strength (**Fig S7B**). Given we did not find sets of CTCF sites with strong preferences for dot versus boundary formation, we conclude that individual sites have a versatile role in specifying chromatin architecture without feature-specific specialization.

**Figure 7.**
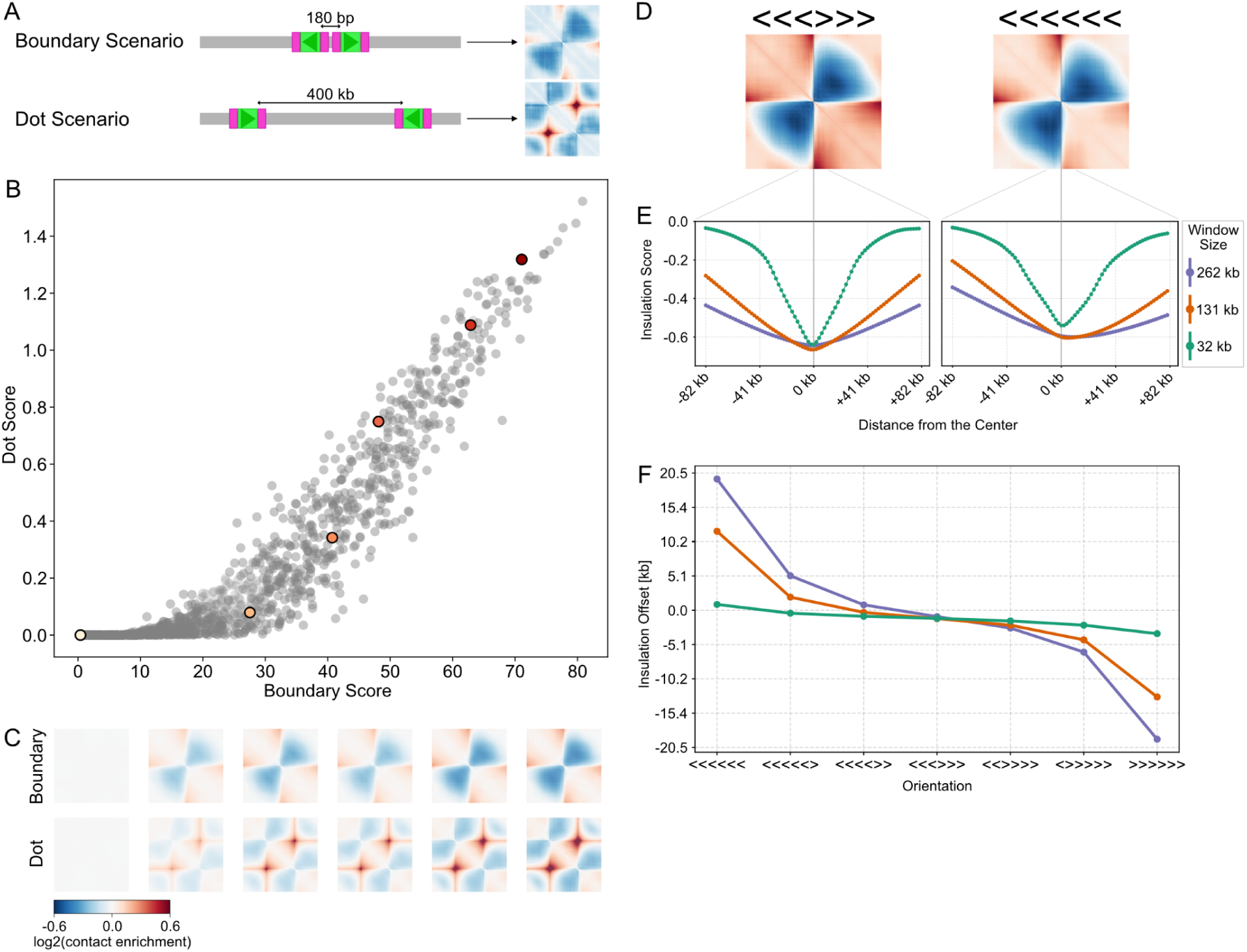
CTCF sites do not mediate feature-specific genome folding. **A)** Illustration of the test for CTCF feature specialization using two distinct layouts: (i) a ‘boundary’ with two divergent sites, 180bp apart, versus (ii) a ‘dot’ with two convergent sites, 400kb apart. CTCF insertions are shown as green rectangles (core motifs) with pink flanks (30bp), arrows indicate the orientation of the CTCF motif. **B)** Scatterplot of boundary vs. dot strength (n=1,500 CTCF sites). Boundary strength is the overall intensity of the map; dot strength is the local average signal within versus around the dot. Six CTCF sites spanning a range of strengths are highlighted with colored dots. **C)** Predicted maps for boundary and dot scenarios for the six highlighted CTCF sites in panel B. **D)** Predicted maps for symmetric (<<<>>>) and asymmetric (<<<<<<) insertions for a cassette of six CTCF sites into the middle of a background sequence. **E)** Insulation scores calculated using sliding diamond windows of three sizes (32.7kb, 131kb, 262.1kb), shown for the central 163,84kb of the map. Note that insulation minima display an offset for the asymmetric case, and the same coloring used for window sizes in E and F. **F)** Insulation minima offset for indicated CTCF cassette insertions. Insulation offset is the position of the insulation score minima relative to the center of the sequence (window sizes of 32.7kb, 131kb, 262.1kb). Each point represents the average across 100 strong CTCF site insertions. Note the insulation offset increases with the asymmetry of the inserted CTCF site configuration and is more pronounced for larger window sizes.

For two CTCF sites inserted in tandem, we observed a slight asymmetry in the predicted maps of co-oriented sites (**Fig 6D, Fig S4D**). When we inserted an asymmetric cassette of six strong motifs, we observed even more asymmetry (**Fig 7D**). To investigate this quantitatively, we examined the insulation profile of the resulting contact maps. To our surprise, for this cluster of six co-oriented motifs, the insulation minima did not align with the center of the inserted cluster, and the misalignment increased with larger insulation window sizes (**Fig 7E**). By inserting cassettes with various orientations of the six sites, we found that both the direction and magnitude of the insulation offset depended on the asymmetry of the inserted cluster (**Fig 7F**). In summary, our analysis revealed that the orientation of CTCF site insertions significantly affects the asymmetry and insulation properties of predicted maps.

## Discussion

In summary, we utilized an updated version of the Akita deep neural network to screen millions of *in silico* DNA sequence perturbations for their impact on genome folding. We quantified the predicted sequence preferences and grammar of CTCF sites with insertion and disruption scores.

For individual CTCF sites, we found a surprisingly low correlation between CTCF ChIP-seq data and *in silico* disruption scores. This aligns with recent observations that CTCF ChIP-seq does not correlate with differential insulation activity [15,17]. Similar to [17], we did not observe a strong correlation between the effectiveness of CTCF sites and their resemblance to the core motif or their degree of conservation. Conversely, a strong correlation between disruption scores and RAD21 ChIP-seq suggests that impactful CTCF sites are able to halt cohesin and block loop extrusion. Our results indicate that CTCF ChIP-seq provides little additional predictive value when single-molecule footprinting data is available, emphasizing the value of emerging methods for studying genome folding.

Our analysis highlights the role of 15bp flanking sequences for specifying strong CTCF sites for genome folding. This generalizes and refines experimental observations about the importance of flanking sequence around core CTCF motifs for insulation activity [15,17]. Although lower information content than the core motif, strong sites displayed sequence preferences both upstream and downstream of the core. While not pronounced enough to be readily extracted by motif-discovery algorithms, the flanking sequences identified by AkitaV2 show similarities to those reported as important for CTCF binding and DNA accessibility [35–40,50]. Similarly to us, [39] and [38] used neural network models to extract sequence preferences around CTCF sites, albeit starting from predicted DNA accessibility instead of predicted genome folding. Differences with [15] likely come from limitations to the number of sites that could be experimentally assayed. [51] reported that accessible sites bound by CTCF can be differentiated from unbound sites by the enrichment of transcription factor binding sites in close proximity to the CTCF motif. However, if co-binding does occur, our results argue for a variable and versatile set of transcription factors.

Our observations argue for two principles of CTCF multi-motif grammar: (i) boundary strength is influenced by the number of CTCF sites within a cluster, and (ii) pattern formation is determined by site orientation and spacing, without intrinsic specialization of CTCF motifs. The first principle aligns with positive correlations between the number of sites and TAD boundary strengths observed in genomic data [45,46], as well as the number of sites and impact on transcription in synthetic sequences [15,16]. The second principle aligns with the correspondence between convergently oriented sites and dots [4] versus divergently oriented sites and boundaries [10] in the genome, as well as orientation-specific impacts on transcription in synthetic sequences [16]. Indeed, we predict that CTCF sites do not preferentially specify dots or boundaries. Experimentally, this is supported by the emergence of new dots between pairs of loci that previously displayed boundaries after the deletion of intervening CTCF sites [7,9]. More broadly, we predict that specific contact patterns emerge from the positions and arrangements of CTCF sites rather than specialized pattern-specific motifs.

A central limitation of our approach is that the sequence preferences and grammar we can identify must have been extracted by the deep neural network we use, AkitaV2. Model performance, architecture, and training scheme could each contribute to what can be learned via our approach [23,52]. The size of the input sequence restricts the maximum distance over which any grammar can be extracted. Only patterns that occur with sufficient regularity have a chance of being learned by the model. For example, repetitive elements that are rare or make species-specific contributions to genome folding are unlikely to be reliably extracted by our approach. Similarly, because large clusters of CTCF sites are relatively rare in the genome (e.g., only 0.36% of boundaries have >10 CTCF motifs), our approach may under-estimate when insulation would saturate as a function of the number of sites in a boundary. Finally, while we primarily used feature-agnostic scores over the full predicted megabase region, feature-specific scores [29] could enable extraction of additional insight.

Collectively, our observations provide a roadmap for the design of experiments hoping to test the sequence determinants of genome folding and downstream consequences for communication between enhancers and promoters. Successful experimental designs will consider both the content of core and flanking CTCF sequences as well as their positioning relative to other regulatory sequences.

## Methods

### Data preprocessing

We followed the preprocessing described in prior research using the Akita framework [19] for the 6 mouse and 5 human datasets in **Supplemental Table 1.** Briefly, we reprocessed these datasets using the distiller pipeline (https://github.com/open2c/distiller-nf, [56]), extracting contacts with pairtools [57], binning each dataset to 2,048bp cooler files (https://github.com/open2c/cooler, [58]) and performing genome-wide iterative correction [59]. Individual target matrices were extracted from genome-wide cooler files for regions corresponding to 1,310,720bp of input sequence, 25% larger than the original 1,048,576bp. As previously, the following steps were applied to matrices for individual regions in the training and test sets: adaptive coarse-graining, normalization for distance-dependence, natural log, clipping to (−2,2), linear interpolation of missing bins, and convolving with a small 2D gaussian filter. The first and third steps used cooltools (https://github.com/open2c/cooltools, [60]).

### Cross-species model

We used the same neural network structure and weights as described for Akita, with the following modification to the last layer: instead of a single dense layer, either a 5 unit dense layer was appended for predicting the 5 human targets or a separate 6 unit dense layer was appended for predicting the 6 mouse targets. We implemented this model using TensorFlow [61]. See https://github.com/calico/basenji/tree/master/manuscripts/akita/v2 params.json for full specification of model weights, learning rate, and other hyperparameters.

### Cross-species training

As for cross-species Basenji training [26], we aimed to avoid leakage between training and test sets by jointly assigning orthologous human and mouse sequences to the same training, validation, or test fold. Briefly this involved: dividing the genome in 5 Mb regions, constructing a bipartite graph if they have >500kb of aligning sequence, and partitioning connected components into 8 distinct folds. We trained an ensemble of models, in which model *i* used fold *i* as its test set, *i*+1 as its validation set, and the remaining folds as its training set. During training, we alternated between batches of 2 human and 2 mouse sequences and Hi-C targets, updating weights for the corresponding final mouse or human dense layer. We trained using stochastic gradient descent with 0.98 momentum and the 1cycle learning rate schedule, in which the learning rate linearly increases from an initial value 0.002 to a maximum value 0.04 over 56 epochs, followed by a linearly decrease back to 0.002 over the next 56 epochs, concluded by dropping the learning rate to 0.0003 for 2 final epochs. We chose the final model weights from the epoch where the validation Pearson’s R reached its max.

### Mouse CTCF sites

To obtain a set of CTCF sites capable of strongly impacting genome folding we took the following steps: extract Jaspar [32] mm10 CTCF site positions (MA0139.1) that overlap with TAD boundaries from [31] obtained at 10kb resolution. These sites were then filtered to exclude overlaps with other CTCF sites (+/- 60bp) or repeat elements (+/- 20bp, RMSK table: https://genome.ucsc.edu/cgi-bin/hgTables?db=mm10&hgta_table=rmsk), resulting in 7,560 CTCF sites.

### Visualizing predicted maps

For visualization of predicted insertions, maps were averaged over ten background sequences and all mouse outputs for the first model, unless specified.

### Predicted Map Signal Strength

Predicted maps were squared, summed, and a square root was taken to yield a positive “Signal Strength” value.

### Disruption Score

The disruption score is calculated as follows: compute the difference between the predicted map before disruption and the predicted map after disruption, square the difference, sum the values, and take the square root. This positive value, averaged across all six cell types and four models unless noted, quantifies disruption.

### Insertion Score

Insertion scores are calculated similarly to disruption scores, where the difference is taken with the background map prediction before insertion. Averaged across six cell types, four models, and ten background sequences per model unless noted.

### Background Generation

We generated backgrounds for each model by iteratively shuffling genomic sequences until the resulting maps achieved a uniformly flat profile, assessed by the predicted map signal strength. Each sequence was shuffled repeatedly until its signal strength fell below a predetermined threshold of 35. Sequences that exceeded this threshold after a maximum of 20 iterations were discarded.

### Dot and Boundary Scores

The boundary score is determined by the global insertion score from a ‘boundary’ scenario insertion. Conversely, the dot score is a localized measure calculated by applying a 13×13 bin kernel to a map patch where a dot is expected. This kernel features a 3×3 bin center over the anticipated dot location, surrounded by four 10×3 bin arms with expected lower signal strength. The dot score is derived from the difference in the average square root of the sum of squared values between the center and the arms of the kernel.

### Dot Anchors

We identified CTCF sites overlapping dot anchors using mm10 ESC data at 10kb resolution [31] using *MUSTACHE* [49], similarly to how we identified those overlapping boundaries. We initially found 39,226 CTCF sites overlapping dot anchors and refined this by excluding 2,278 sites that also overlapped TAD boundaries, resulting in a distinct set of 36,948 CTCF sites.

### Insulation Offset

The insulation score is derived from the average value within a sliding diamond window along the main diagonal of the map, similar to the method used in [60]. The insulation offset is the distance between the insulation score minima and the map’s center point. This offset was averaged across the tested set of CTCF sites, all cell types, ten different background sequences, and four distinct models.

### Statistics and software

The neural network has been implemented using python (v3.7) and tensorflow (v2.4). The main text and figure legends indicate the statistical tests used in the comparisons. Pearson R and Spearman R were calculated with scipy 1.11.4 [62]. Analyses were performed using: numpy 1.23.5 [63], pandas 2.1.4 [64], matplotlib 3.8.4 [65], and seaborn 0.13.0 [66], and additionally made use of h5py 3.10.0 [67], and pysam 0.22.0 [68].

### Code availability

Scripts used for cross-species AkitaV2 training and model weights are available at https://github.com/calico/basenji/tree/master/manuscripts/akita/v2. General utilities for AkitaV2 are available at https://github.com/Fudenberg-Research-Group/akita_utils. Code to reproduce analyses, including figures, are available at https://github.com/Fudenberg-Research-Group/akitaV2-analyses.

## Acknowledgements

The authors thank Elphège Nora, Erika Anderson, and Katherine Pollard for feedback. G.F. and P.N.S. are supported by the National Institute of General Medical Sciences R35 GM143116-01.

## Supplementary Table

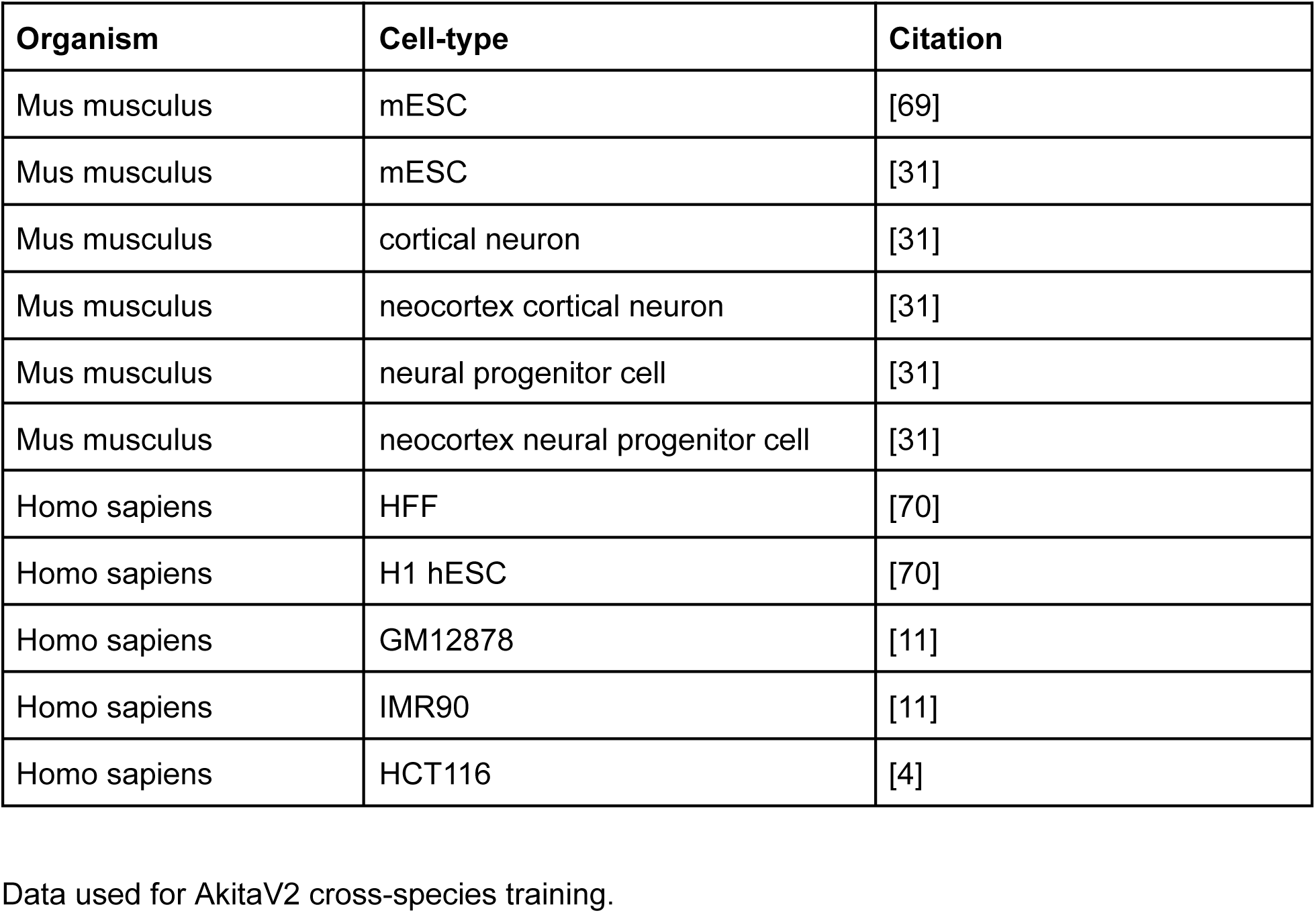

## Supplementary Figure

**Supplementary Figure 1.**
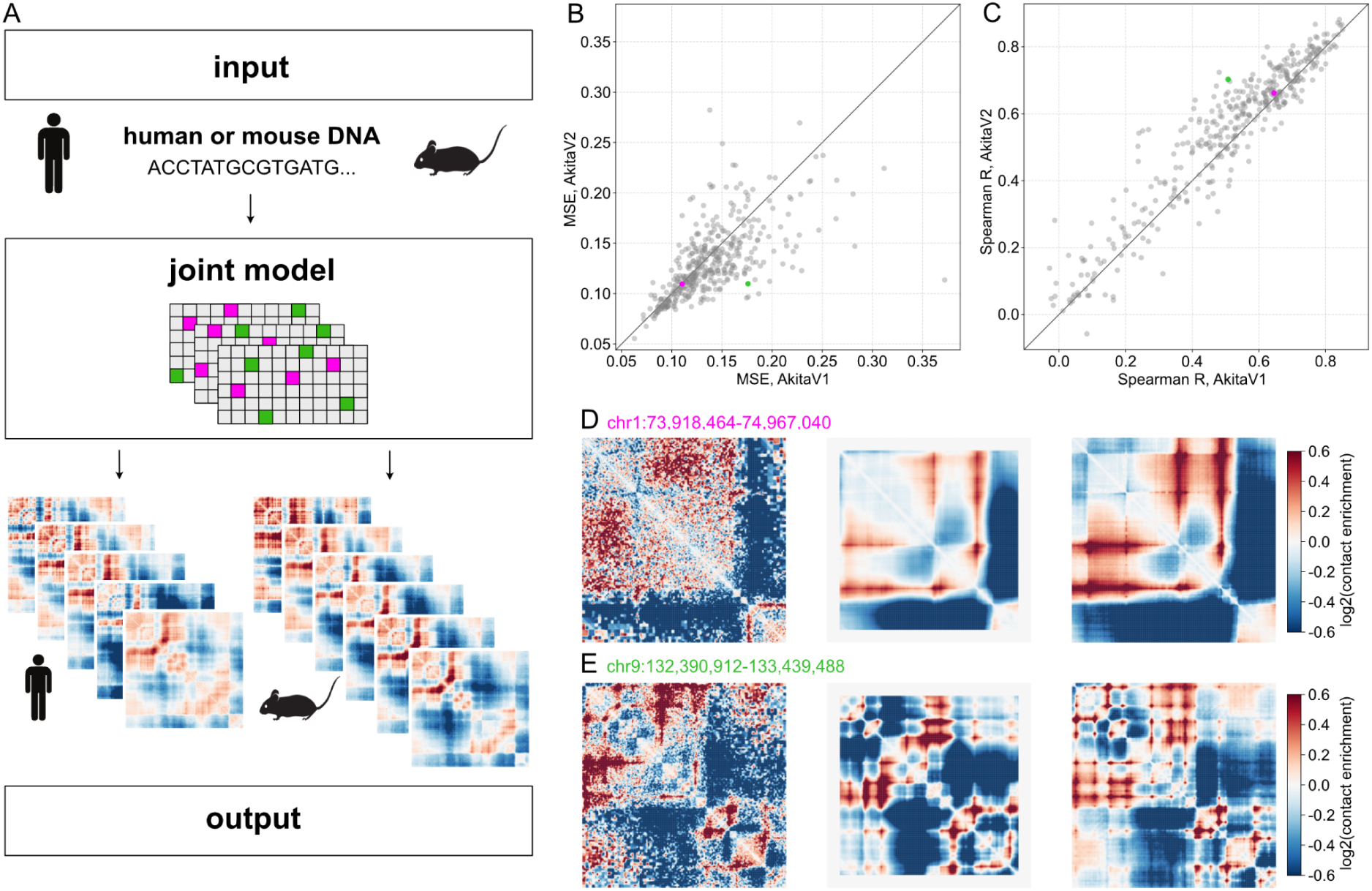
AkitaV2 enables mouse and human predictions via a cross-species training approach. **A)** AkitaV2 architecture. This model inputs ∼1.3 million base pairs of DNA to predict log(observed/expected) pairwise contact frequencies. The model employs a shared trunk and two distinct prediction heads: one for six mouse cell types and another for five human cell types. **B)** Scatterplot of MSE for AkitaV1 vs. AkitaV2 for each genomic region in the AkitaV1 test set. AkitaV2 displays enhanced performance (0.131 vs. 0.139). Pink and green dots highlight two representative genomic regions: one where AkitaV2 prediction is comparable with AkitaV1, and one where the AkitaV2 prediction outperformed AkitaV1. Predicted maps for the two highlighted regions are shown below. **C)** Scatterplot of Spearman correlation coefficients for AkitaV1 vs AkitaV2 for regions in the test set. Akita displayed improved Spearman R (0.59 vs. 0.56) and Pearson R (0.66 vs. 0.62) across the test set. Colored dots as in (B). **D)** Visual comparison of log(observed/expected) contact frequencies for a genomic window with minimal improvement. From left to right: the experimental target map, the prediction by AkitaV1, and the prediction by AkitaV2. **E)** Comparison of log(observed/expected) contact frequencies for a genomic window with visible improvement. From left to right: the experimental target map, followed by predictions from AkitaV1 and then by AkitaV2.

**Supplementary Figure 2.**
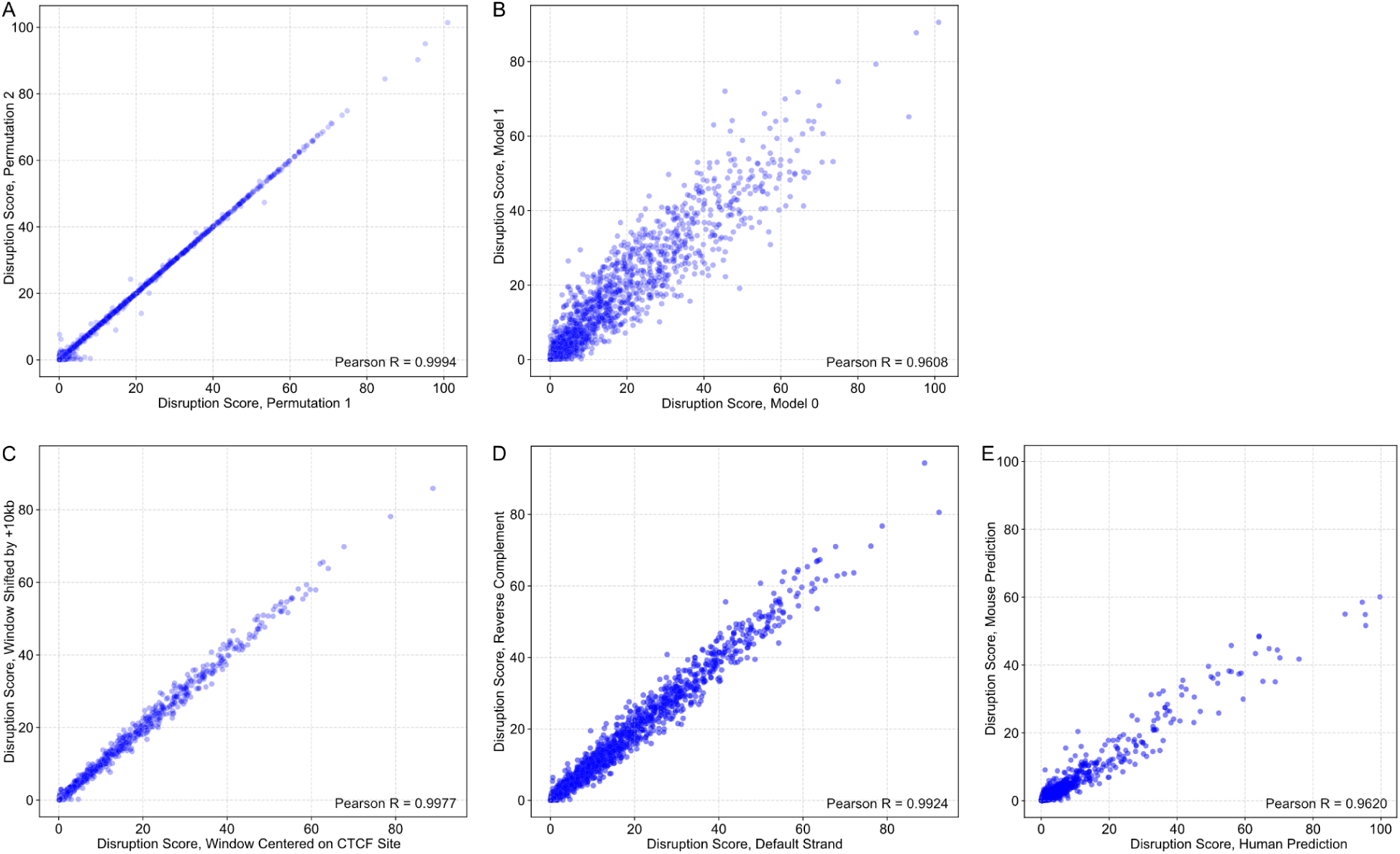
DNA sequence disruption by permutation offers a robust strategy for computing predicted impacts of CTCF. **A)** Disruption scores are highly correlated for random CTCF permutations. Scatterplot of disruption scores for n=7,560 individual CTCF sites subjected to random motif permutations, where each point represents the predicted disruption score of an individual CTCF site. Disruption scores were computed twice (with model 0) for each CTCF-binding site overlapping a TAD boundary. **B)** Inter-model consistency in CTCF site disruption. Scatterplot of disruption scores for n=7,560 individual CTCF sites compared between model 0 and model 1. Disruption scores are highly correlated across all pairs of models 0-7 (PearsonR > 0.955). **C)** Position of disrupted CTCF site relative to the prediction window. This plot explores the effect of shifting the predictive window by +10kb. We tested shifts of ±10kb, ±1kb, ±100bp, ±10bp, and ±1bp, comparing the scores for each shift with the scores from the centered permutation. All correlation coefficients between the disruption scores for shifted permutations and the centered permutation were consistently high, exceeding 0.997. **D)** Consistency in CTCF sites disruption across DNA strands. Disruption scores do not depend on the input sequence orientation. **E)** Inter-species consistency of CTCF site disruption. 18Mb of mouse chromosome 1 (ch1:3,653,632-21,776,376) was disrupted by permuting a sliding ∼200bp window and disruption scores were calculated for either the human (hESC) or mouse (mESC) output. Scatterplot shows median disruption score across four models for each 200bp window for the human versus the mouse output (Pearson R > 0.96).

**Supplementary Figure 3.**
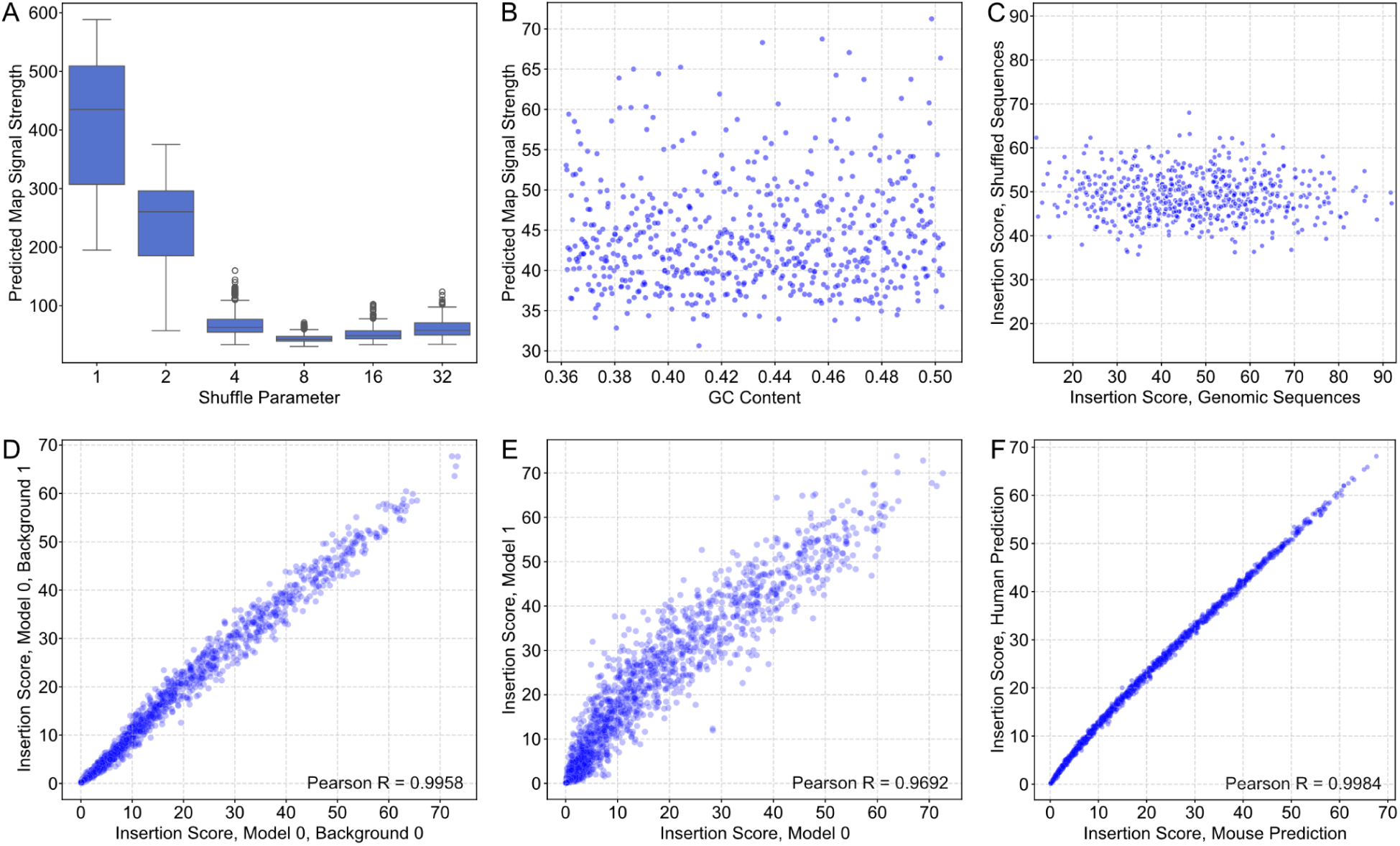
DNA Sequence Shuffling and Insertion Score Robustness. **A)** Boxplots constructed from predicted map signal strength scores of n=590 shuffled genomic windows show DNA sequence shuffling impact on contact matrix strength, with predicted map signal strength for model 0 across shuffled sequences (1, 2, 4, 8, 16, 32 nucleotides). Lower scores denote weaker maps, with k=8 shuffling resulting in the most neutral maps. **B)** Scatter plot of predicted map signal strength versus GC content for shuffled genomic sequences. Points represent scores from model 0 for n=590 genomic windows shuffled using k=8, and show no trend between GC content and SCD. **C)** Scatterplot comparing insertion scores for the insertion of a strong CTCF site into n=590 genomic sequences, both original and shuffled once with k=8. While shuffling does not alter the mean, it remarkably reduces the variance of the insertion scores. **D)** Scatter plot of virtual insertion score for background sequence 0 vs. background sequence 1 for model 0 across n=7560 CTCF sites. Virtual insertion scores are highly correlated across backgrounds (PearsonR > 0.94 for any pair of background sequences). **E)** Scatterplot of insertion scores between model 0 and model 1 for n=7560 CTCF sites. Virtual insertion scores are highly correlated across pairs of models (PearsonR > 0.96). **F)** Scatterplot of insertion scores between mouse and human predictions for n=7560 mouse CTCF sites inserted into shuffled mouse background sequences. Insertion scores are highly correlated (PearsonR > 0.99).

**Supplementary Figure 4.**
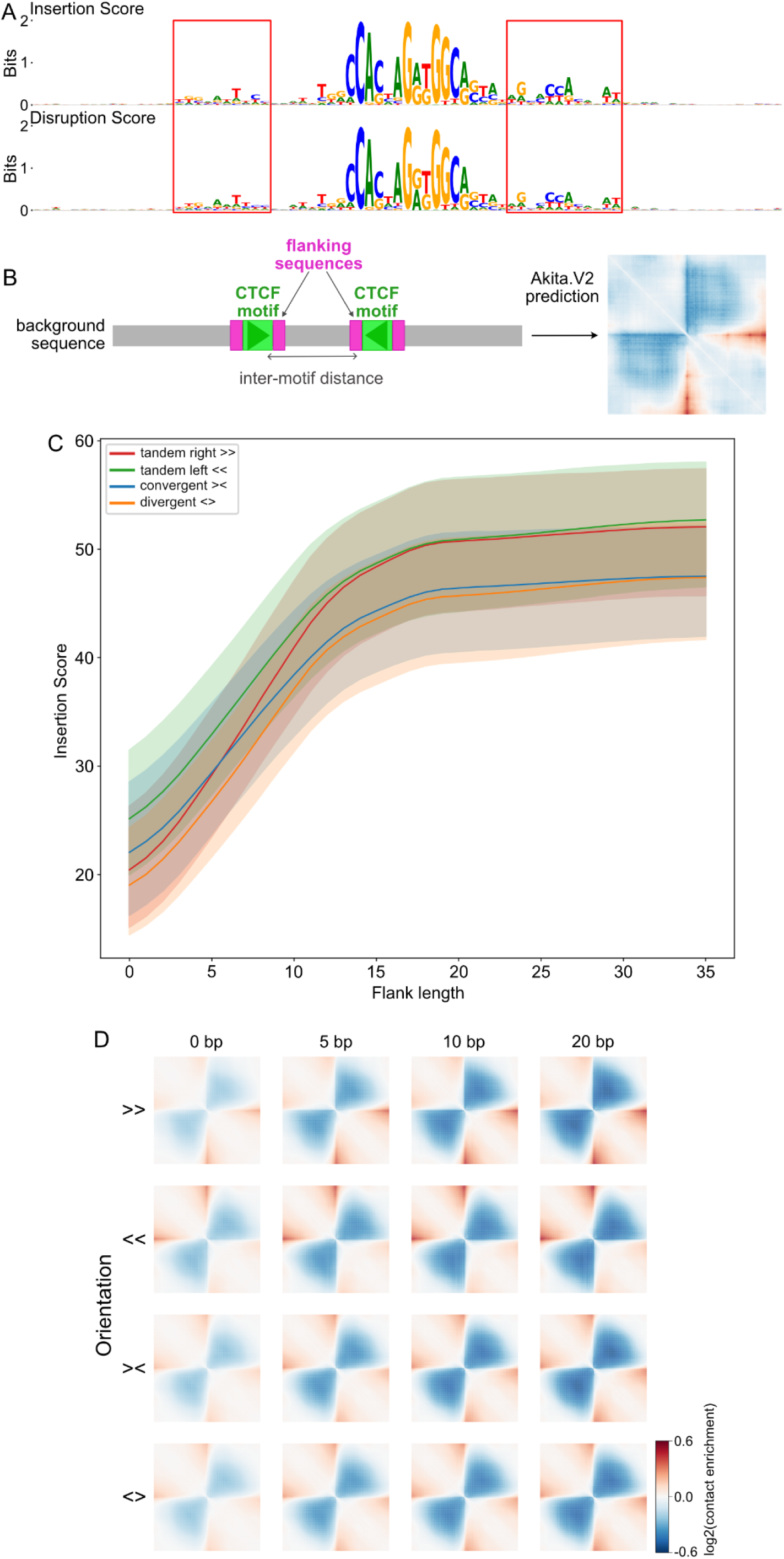
Sequence Preferences within Flanking Sequences and the Double-Site Virtual Insertion. **A)** Sequence logos of CTCF motifs and 30bp flanking sequences from the strongest 150 motifs ranked by either insertion (top) or disruption (bottom) scores.Red boxes highlight weak sequence preferences upstream and downstream of the CTCF motifs. **B)** Illustration of a double CTCF site insertion. Two CTCF sites (green boxes) are virtually inserted symmetrically around a background sequence’s midpoint (gray rectangle) with constant 180bp inter-motif distance (as in [15]) with flanking regions (pink boxes). The impact is quantified by the squared contact difference between maps with and without inserted CTCF sites (insertion score). **C)** Insertion score versus flanking sequence length for insertions of tandem CTCF sites in four orientations (left, right, convergent, divergent). Grouping and shading as in **Fig 6B**. Insertions of tandem sites show a similar trend to single sites, and similar trends across all orientations. **D)** Predicted maps for double CTCF insertions with increasing flank length for different orientations. Inserted CTCF sequence extracted from chr7:37,357,852-37,357,871. The maps are arranged in a grid by flank length (columns) and site orientation (left, right, convergent, divergent) in rows, demonstrating similar strength impacts across orientations but slight asymmetry for sites inserted in tandem.

**Supplementary Figure 5.**
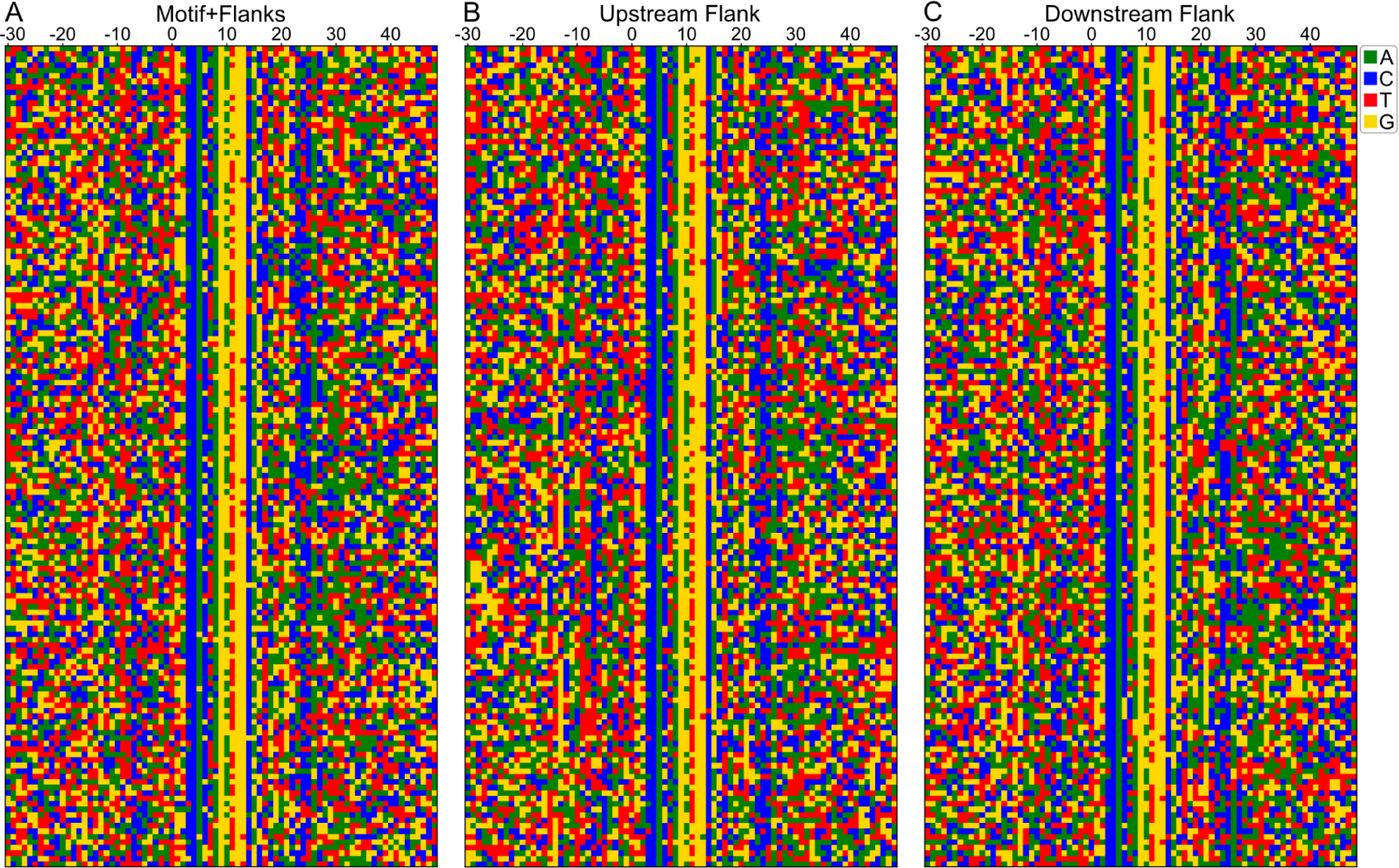
No evidence of clear clusters in flanking sequences. A) Heatmap of nucleotide composition around 150 strong CTCF sites (±30bp), with rows sorted according to the Hamming distance of motifs combined with their flanking sequences. B) Same as panel A except rows are ordered by the Hamming distance of the upstream flank. C) Same as panel A and B, but with rows arranged by the Hamming distance of the downstream flank.

**Supplementary Figure 6.**
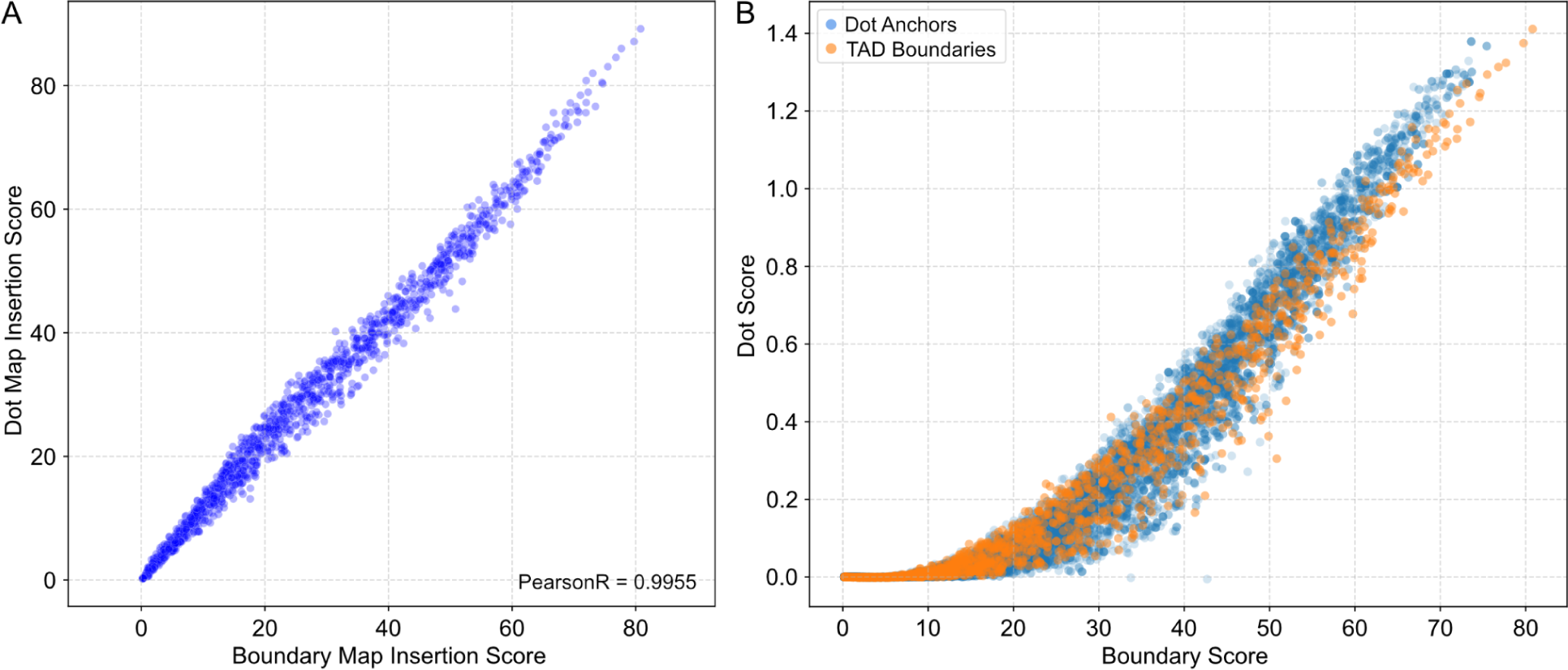
Feature-Specificity of CTCF Sites. **A)** Insertion score correlation between boundary and dot scenarios. A scatter plot comparing insertion scores from boundary to dot scenarios shows a high correlation (PearsonR > 0.99), indicating that the global metric of predicted map strength does not significantly vary with the insertion scenario. **B)** CTCF sites’ genomic origin vs. feature-specificity. A scatter plot compares CTCF sites, differentiated by overlap with TAD boundaries (orange dots, n=1,500) or dot anchors (blue dots, n>36,900), against boundary and dot strengths. Both sets of CTCF sites are disjoint. Uniform distribution across the plot shows all sites behave similarly in the experiment, regardless of their genomic origin. This suggests that CTCF’s role in chromatin architecture does not inherently differ between those overlapping with TAD boundaries and those at dot anchors.

